# Intravital imaging of *Candida albicans* identifies differential in vitro and in vivo filamentation phenotypes for transcription factor deletion mutants

**DOI:** 10.1101/2021.05.10.443530

**Authors:** Rohan S. Wakade, Manning Huang, Aaron P. Mitchell, Melanie Wellington, Damian J. Krysan

## Abstract

*Candida albicans* is an important cause of human fungal infections. A widely studied virulence trait of *C. albicans* is its ability to undergo filamentation to hyphae and pseudohyphae. Although yeast, pseudohyphae and hyphae are present in pathological samples of infected mammalian tissue, it has been challenging to characterize the role of regulatory networks and specific genes during in vivo filamentation. In addition, the phenotypic heterogeneity of *C. albicans* clinical isolates is becoming increasingly recognized and correlating this heterogeneity with pathogenesis remains an important goal. Here, we describe the use of an intravital imaging approach to characterize *C. albicans* filamentation in a mammalian model of infection by taking advantage of the translucence of mouse pinna (ears). Using this model, we have found that the in vitro and in vivo filamentation phenotypes of different *C. albicans* isolates can vary significantly, particularly when in vivo filamentation is compared to solid agar-based assays. We also show that the well-characterized transcriptional regulators Efg1 and Brg1 appear to play important roles both in vivo and in vitro. In contrast, Ume6 is much more important in vitro than in vivo. Finally, strains that are dependent on Bcr1 for in vitro filamentation are able to form filaments in vivo. This intravital imaging approach provides a new approach to the systematic characterization of this important virulence trait during mammalian infection. Our initial studies provide support for the notion that the regulation and initiation of *C. albicans* filamentation in vivo is distinct from in vitro induction.

**Importance:** *Candida albicans* is one of the most common causes of fungal infections in humans. *C. albicans* undergoes a transition from a round yeast form to a filamentous form during infection which is critical for its ability to cause disease. Although this transition has been studied in the laboratory for years, methods to do so in an animal model of infection have been limited. We have developed a microscopy method to visualize fluorescently labeled C. albicans undergoing this transition in the subcutaneous tissue of mice. Our studies indicate that the regulation of C. albicans filamentation during infection is distinct from that observed in laboratory conditions.

## Introduction

Microbial virulence traits and factors are frequently studied using in vitro experimental systems, particularly when the goal is to probe detailed molecular mechanisms of pathogenesis. The premise of such experiments is based on a correlation between the in vitro observations and the events that occur during infection of the host; this assumption is frequently quite reasonable but also can be experimentally challenging to verify. Here, we describe the use of a novel intravital imaging approach to characterize the in vivo ability of *Candida albicans* to transition from yeast to filamentous morphology, a key virulence trait in this important, highly prevalent human fungal pathogen (1).

*Candida albicans* is one of the most common human fungal pathogens and causes both superficial mucosal infections as well as invasive infections of organs such as liver, spleen, kidney and brain. *C. albicans* undergoes characteristic morphologic transitions between round yeast and filamentous hyphae and pseudohyphae (2). Histopathologic analyses indicate that all three morphologic forms of *C. albicans* are generally present within infected anatomic sites. The transcriptional regulation of *C. albicans* filamentation has been the subject of extensive study and has led to the identification of TFs that play roles in this morphogenetic transition (3, 4). Based on the study of three key hyphae-associated TFs (*EFG1, BRG1*, and *UME6* along with the biofilm regulator Bcr1) in the standard reference strain SC5314 and four different clinical isolates of *C. albicans*, Huang et al. found that the transcriptional circuitry regulating in vitro biofilm formation and filamentation varied significantly among the strains (5).

We were interested in determining the roles of these TFs during in vivo filamentation. It is clear from a variety of studies that the ability of a given *C. albicans* mutant to undergo filamentation in vitro can vary with the specific in vitro inducing stimulus (6, 7). The existence of condition-dependent filamentation programs was nicely demonstrated by the systematic analysis reported by Azadmanesh et al (6). We hypothesized that filamentation during mammalian infection may have characteristics that are distinct from in vitro filamentation. Currently, there are limited approaches to directly studying *C. albicans* morphologic transitions during infection. Histologic analyses of infected organs can provide information about filamentation. However, quantitative analysis is difficult because hyphae sectioned perpendicular to the long axis can appear as yeast. The zebrafish model has recently been used to advantage to characterize filamentation in vivo and provided a number of insights into the roles of both yeast and filaments during infection (8). For mammalian models, Witchley et al. recently reported a FISH-based approach that is applicable to the quantitative characterization of *C. albicans* filamentation during the colonization of the murine GI tract (9).

Here, we report a novel intravital microscopy approach that has allowed us to characterize the *C. albicans* yeast-to-filament transition in a mouse model of infection (10). *C. albicans* is both a commensal colonizer of the human gastrointestinal (GI) tract and a cause of invasive infections (1). A well-accepted model for the transition from commensal colonization to pathogenic dissemination (11) begins with *C. albicans* breeching the epithelial cell layer of a mucosal tissue such as the oral cavity or the GI tract to invade the subdermal/submucosal stroma (Fig. 1A). Next, the fungus gains access to the vascular system by traversing the endothelial cells of blood vessels and, ultimately, disseminates to target organs such as the kidney, liver, spleen and brain. Interactions of *C. albicans* with epithelial cells, endothelium, and target organs have been studied extensively but little is known of its interactions with subepithelial tissue (12). To study this stage of infection and to characterize in vivo filamentation of *C. albicans*, we adapted an intravital imaging method developed in our lab in which fluorescently labeled *C. albicans* are directly injected into the ear of mice (10) and observed using confocal microscopy (Fig. 1B). Using this approach, we demonstrate that: 1) the correlation between in vitro and in vivo filamentation phenotypes is dependent on the specific in vitro induction stimuli and 2) the transcriptional regulation of in vivo filamentation is distinct from in vitro filamentation.

**Figure 1.**
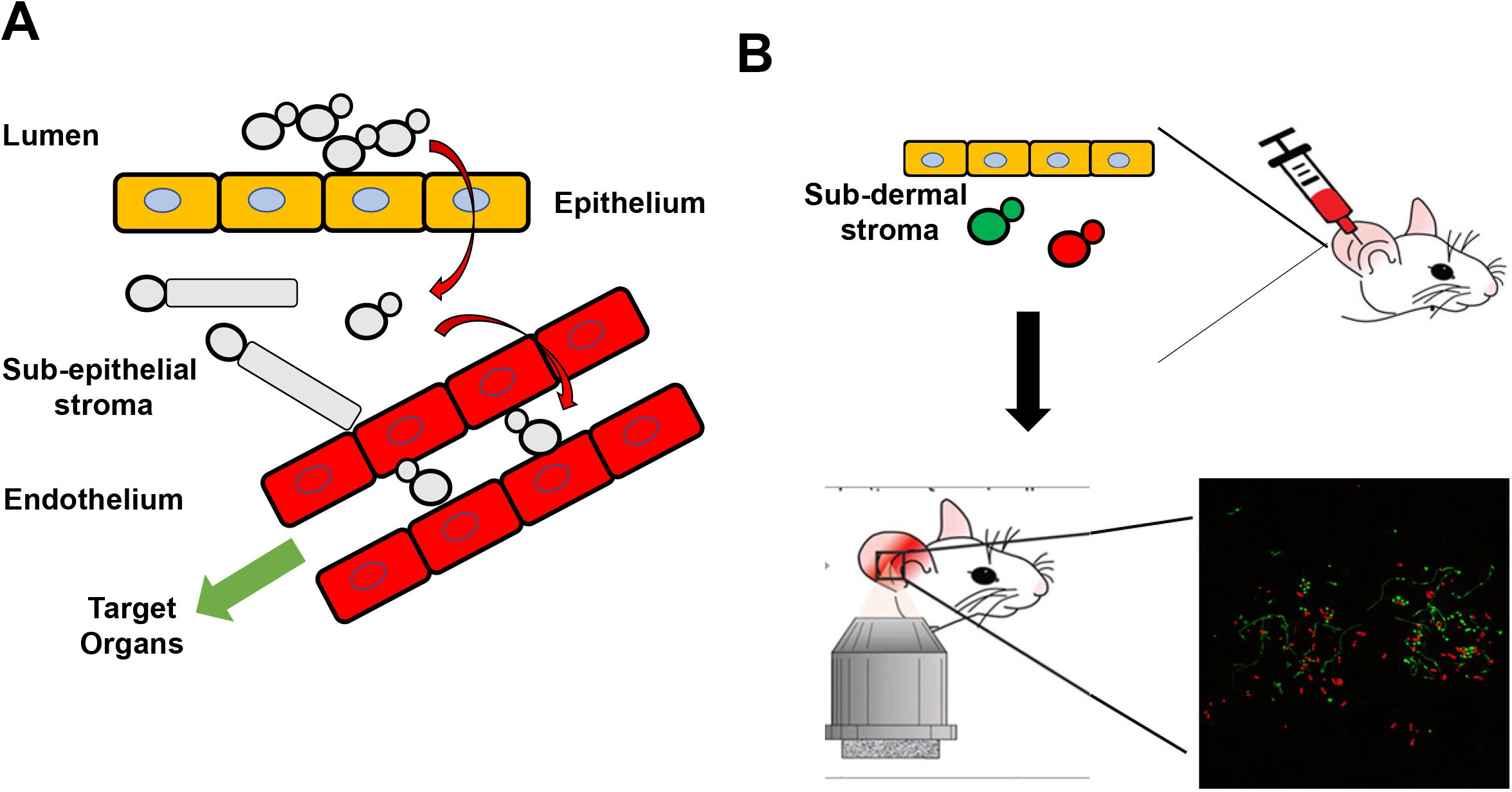
Premise and schematic of intravital imaging of *Candida albicans* in ear tissue of mice. A. Model of translocation and dissemination of *C. albicans*. B. Schematic representing injection of orthogonally-labeled fluorescent *C. albicans* and imaging with confocal microscope.

## Results

### The correlation between in vitro and in vivo filamentation phenotypes varies with the specific in vitro induction stimuli

To characterize the correlation between in vitro and in vivo filamentation phenotypes, we first constructed a GFP-labelled derivative of the standard reference strain SC5314 and injected it into the ears of DBA/2 mice (10); this strain of mice lacks complement C5 which limits initial edema due to reduced influx of phagocytes and thereby improves resolution. SC5314 undergoes robust filamentation in this model at 24 hours postinfection (Fig. 2A). Although we can clearly distinguish yeast cells from filamentous cells (Fig. 2B), we are not able to consistently distinguish between hyphae and pseudohyphae and thus score filaments as not-yeast; see materials and methods section for a complete description of scoring method. We also induced hyphae formation in vitro using RPMI medium supplemented with 10% bovine serum (Fig. 2C) for 4 hours. For SC5314, comparable numbers of filaments are observed at 24 hours in vivo and after 4 hours of in vitro induction (Fig. 2D). This is consistent with the general observations in the literature indicating that SC5314 forms robust filaments under both liquid and plate-based conditions (5, 13).

**Figure 2.**
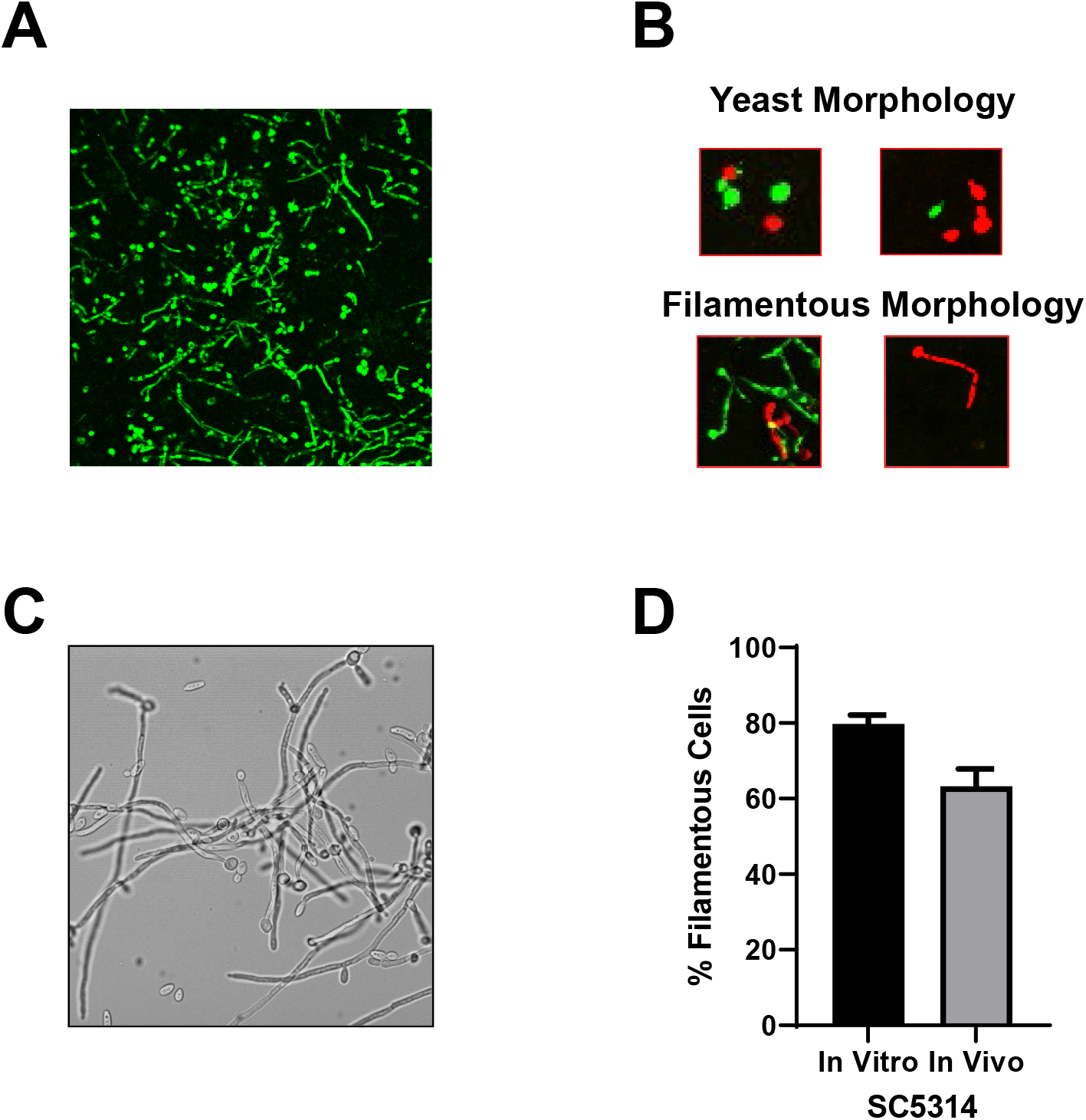
Comparison of filamentation of reference strain SC5314 in vitro and in vivo using the intravital imaging approach. A. Representative field showing GFP-labeled SC5314 within tissue of the ear 24 hr post-infection. B. Examples of yeast and filamentous morphologies as captured by intravital imaging assay. C. SC5314 cells exposed to RPMI+10% serum at 37°C for 4 hr. D. Comparison of percentage of filamentous cells in vitro (RPMI+10% serum at 37°C for 4 hr) and in vivo (24 hr post-infection). Bars indicate mean of 4-5 fields from replicate experiments with error bars indicating standard deviation.

To extend this analysis to strains with heterogenous filamentation phenotypes, we took advantage of the recent characterization of strains from a set of 21 clinical isolates that had also previously been characterized for virulence phenotypes (13, 14). We chose four strains (P87, P57010, P57055, and P76067) from different clades that had relative filamentation scores on solid Spider medium of P87~SC5314~P76067>>P57055~P75010 while in liquid RPMI without serum the relative filamentation was: SC5314~P87>P7607>>P57055~P75010 (5, 12). The strains were fluorescently labelled and assessed both in vitro and in vivo as described for the reference strain SC5314 (Fig. 3A-B). The addition of serum to RPMI medium induced P57055 to filament much more relative to RPMI alone (~5% to 65%) otherwise the relative order of in vitro filamentation phenotypes was similar to RPMI alone. Although the strain with the lowest amount of filamentation in vitro (P75010), formed three-fold more filaments in the presence of serum relative to the absence based on the data from Hirakawa et al. (13).

**Figure 3.**
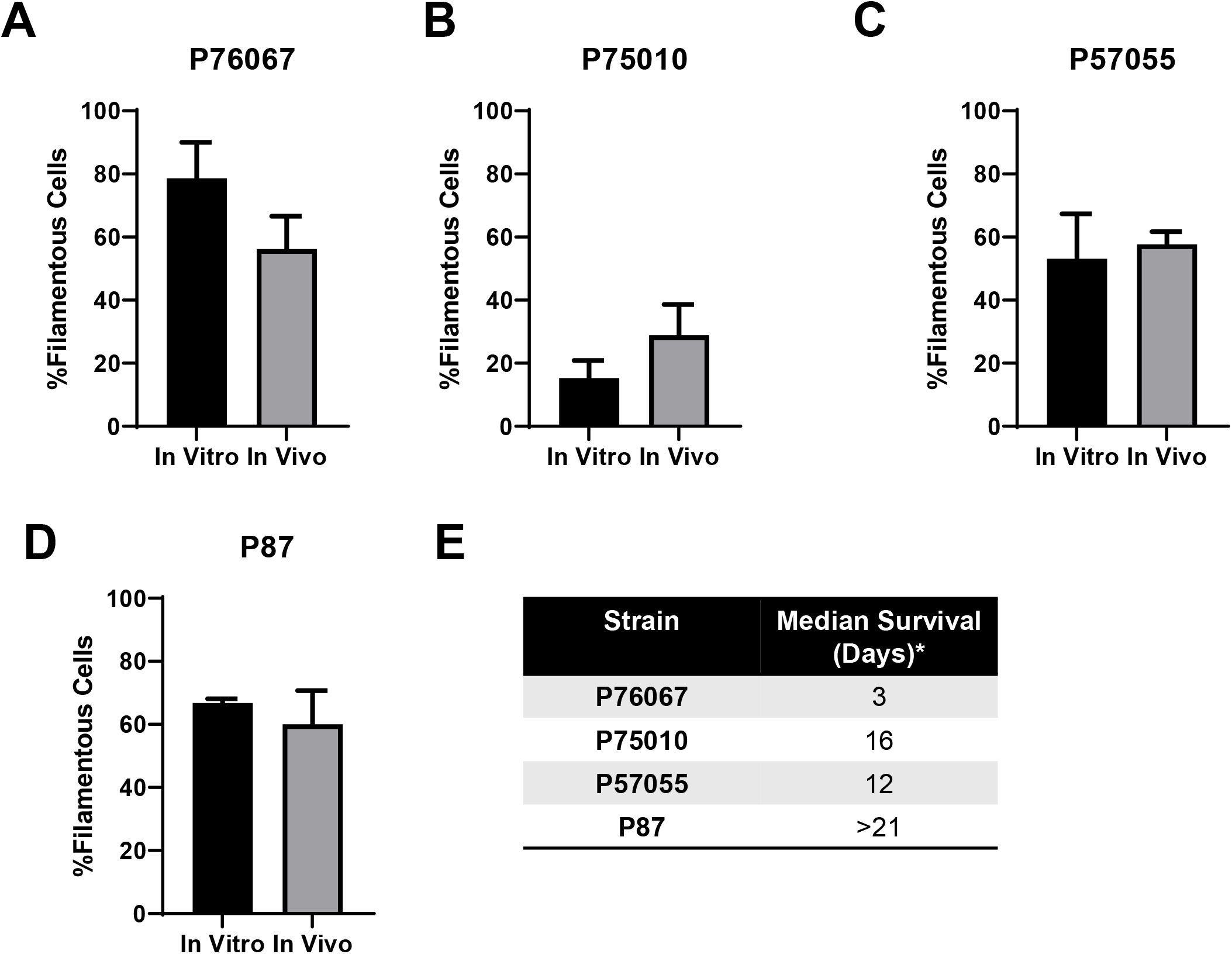
Comparison of in vitro and in vivo filamentation across four well-characterized clinical isolates of *C. albicans*. A-D. The indicated isolates were imaged after exposure to RPMI+10% serum at 37°C for 4 hr (in vitro) or 24 hr post-injection into the ears of mice (in vivo). Bars indicate mean of 4-5 fields from replicate experiments with error bars indicating standard deviation. E. Median survival of the clinical isolates calculated from data reported by Wu et al. (14).

In vivo, three of the strains filamented to similar extents (P87, P76067, and P57055) and matched well with SC5314. Importantly, however, the strain that formed the least amounts of filaments (P75010) only differed by ~2-fold from the other strains. Thus, the overall variation in filamentation phenotypes in vivo was less than observed in vitro, particularly with respect to the filamentation scores on solid medium (13). Indeed, the relative order of filamentation in RPMI+10% serum matched that seen in vivo much better than solid Spider medium and slightly better than RPMI alone (13). Hirakawa et al. (13) and Azadmanesh et al (6) had examined whether in vitro filamentation of these clinical strains or mutants correlated with virulence but had found no clear relationship. The four clinical isolates we examined have very different median survival rates as reported by Wu et al (Ref. 14, Fig. 3E). For example, P87, which filamented well under all three in vitro conditions and in vivo, was the least virulent strain with no definable time to 50% survival. In addition, the variation in the virulence phenotypes reported for the other three strains is much wider than the variation in their relative abilities to filament in vivo. Additional studies of clinical isolates will be needed to establish a well-powered correlation between in vivo filamentation and virulence.

### Validation of a dual fluorophore assay to assess the effect of mutations on *C. albicans* filamentation during infection

The transcriptional regulation of *C. albicans* filamentation has been the subject of extensive study (3, 4, 5, 6, 7) and has led to the identification of a set of transcription factors (TFs) that play a role in filamentation. Previously, we and others have shown that this network of TFs appears to be dependent upon the specific environmental context for the filamentation (6, 7). Therefore, we hypothesized that the transcriptional regulation of filamentation during infection may have distinct patterns relative to in vitro conditions. To test this hypothesis, we applied our intravital imaging assay to the characterization of the ability of different TF deletion strains to undergo filamentation in vivo. In order to directly compare a given mutant to a reference control strain, we infected animals with an inoculum containing a 1:1 ratio of a reference strain (SN background) labeled with GFP and an iRFP-labeled homozygous *EFG1* deletion mutant derived from that reference strain. As shown in Fig. 4A/B, *EFG1* is required for filamentation under both in vitro and in vivo conditions. Next, we examined a strain that is constitutively filamentous in vitro due to deletion of a transcriptional repressor of filamentation, *TUP1* (15). Hyper-filamentous strains are difficult to study using the intravenous inoculation model because they fail to establish infection. Consistent with its in vitro phenotype, only filamentous forms of the *tup1*ΔΔ mutant were observable both in vivo and in vitro (Fig. 4C/D).

**Figure 4.**
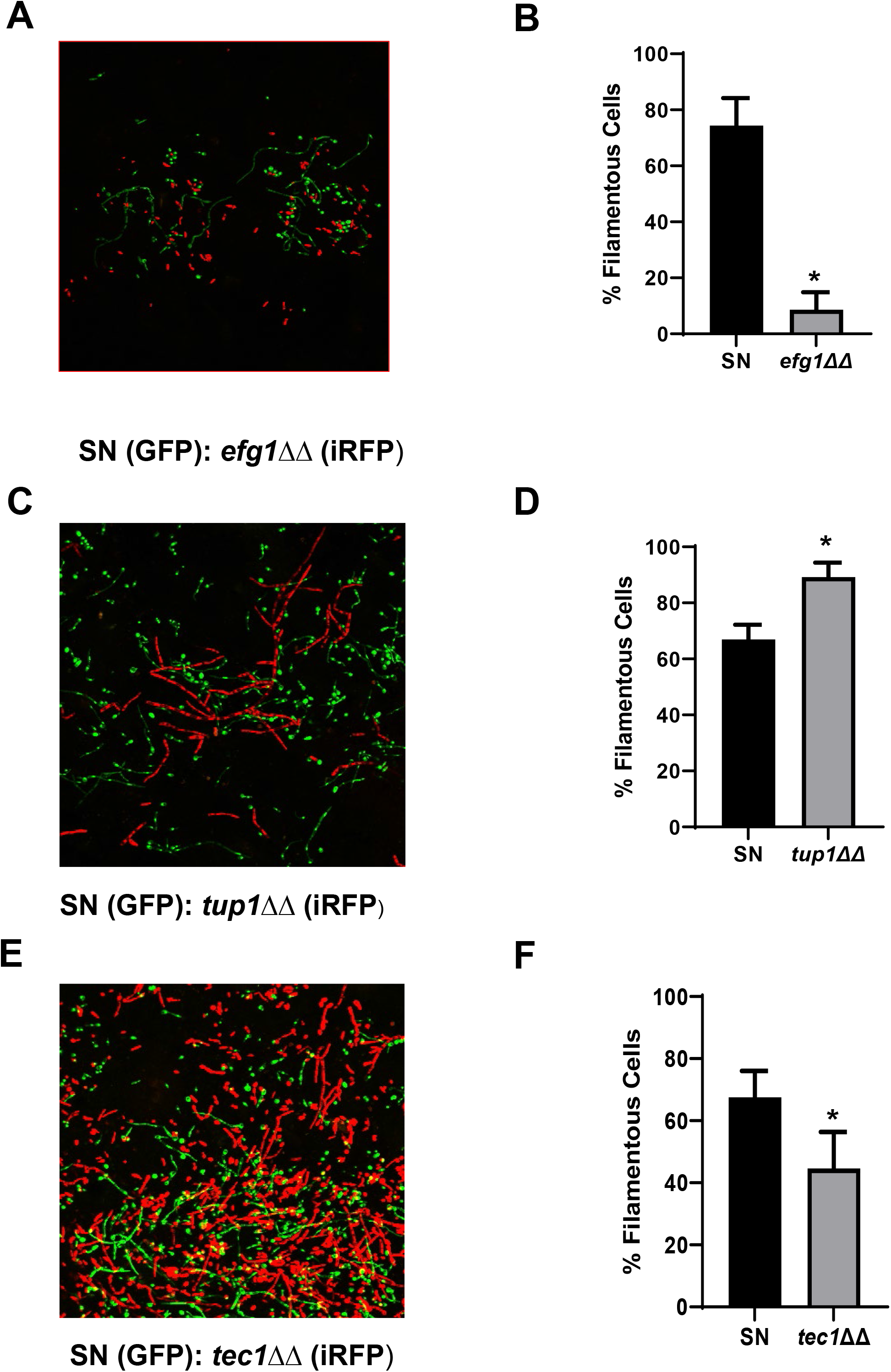
Validation of WT:mutant mixed infection model to assess effect of transcription factor deletion strains on in vivo filamentation. A/C/E. Representative fields for 1:1 WT(GFP):TF deletion mutant (iRFP) infections after 24 hours post-infection with a 1:1 mixture of the indicated strains. B/D/F. Bars indicate mean of 4-5 fields from replicate experiments with error bars indicating standard deviation. * indicates *P* <0.01 for a Student’s t test comparing WT filamentation ratio to the indicated TF mutant.

Finally, we tested the ability of a strain lacking *TEC1* to filament in vivo. Tec1 is regulated by Efg1 and Cph2 in vitro and is required for full virulence (16, 17). Based on histological sections of mouse kidneys infected with a *tec1*ΔΔ strain, it appears that this strain retains the ability to filament in vivo despite being deficient in almost all in vitro conditions reported. As shown in Fig. 4E/F, the *tec1*ΔΔ strain forms filaments in vivo but the ratio of filaments to yeast is reduced relative to the reference strain (*P* = 0.003, Student’s t test). These experiments confirm that the assay can identify both hypo- and hyper-filamentous mutant strains. The discordant phenotype previously reported for in vitro and in vivo filamentation phenotypes for the *tec1*ΔΔ strain is recapitulated in our model and further suggests that inducers of filamentation in the stromal tissue of the ear may be similar to those operative in the kidney.

### Efg1 and Brg1 mutations reduce in vivo filamentation in *C. albicans* clinical isolates

Once we validated the ability of the in vivo imaging assay to characterize mutants with both hypo- and hyper-filamentation phenotypes, we examined the effect of deleting master regulatory TFs in the five strains characterized above (5). Efg1 is one of the most widely studied transcriptional regulators of *C. albicans* and has been shown to be required for filamentation under both in vitro and in vivo conditions (18). Huang et al. found that Efg1 was critical to biofilm formation and in vitro filamentation in all five of the strain backgrounds (5). To extend our finding that it is required for in vivo filamentation in the SN genetic background, we labelled *efg1*ΔΔ mutants in SC5314, P87, P57010, P57055, and P76067 and tested each strain’s ability to filament in our standard in vitro conditions and in vivo. In vitro, deletion of *EFG1* reduced filamentation in all strains except P75010 which formed very few filaments at baseline (Fig. 5A/B); these data were similar to those previously reported by Huang et al. (5). Similarly, *efg1*ΔΔ mutants were significantly impaired for filamentation in vivo with only mutants in the P57050 background forming more than 10% filaments (Fig. 5C/D).

**Figure 5.**
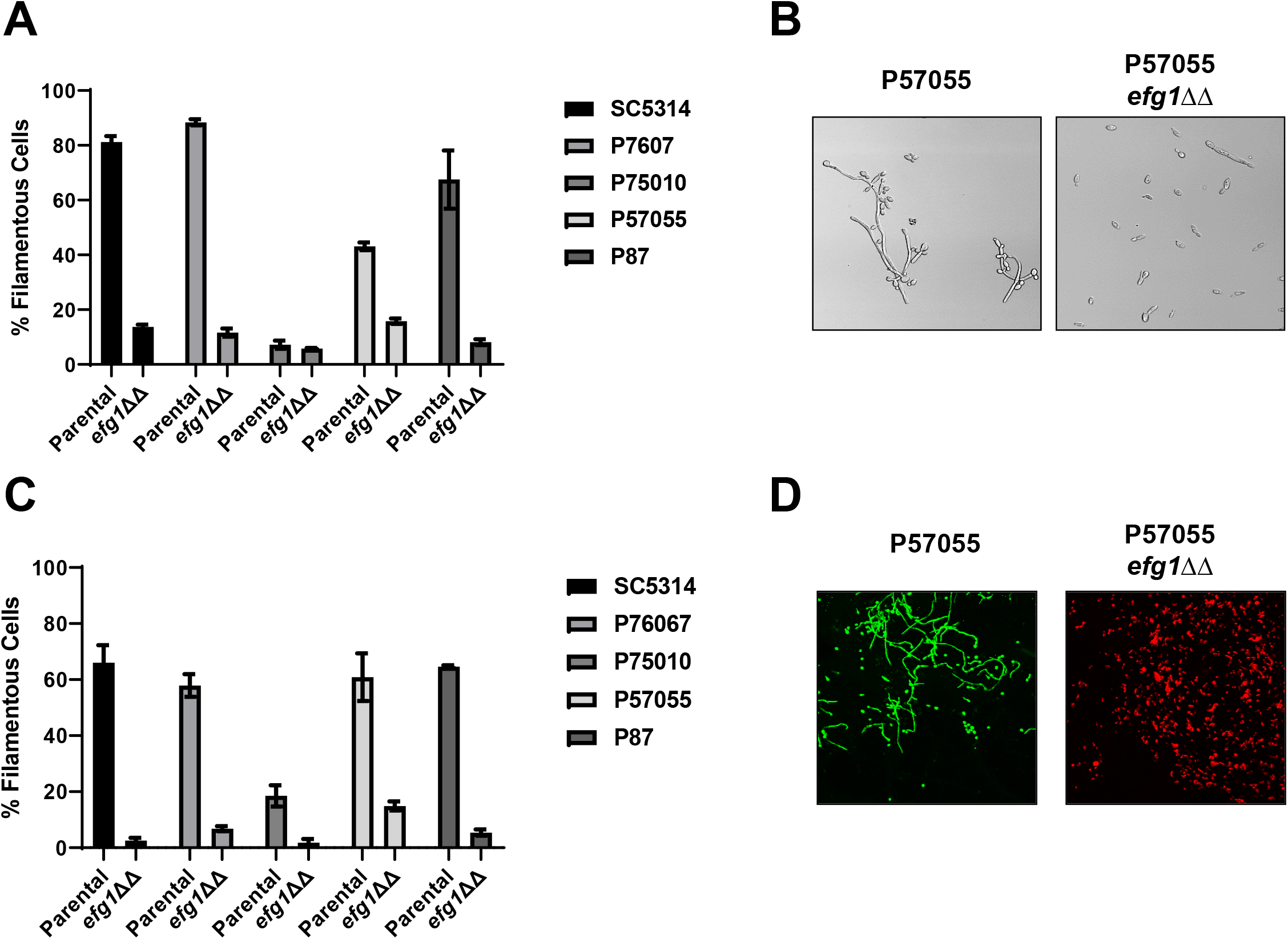
Efg1 is required for in vitro and in vivo filamentation across multiple *C. albicans* isolates. A. Comparison of the in vitro filamentation (RPMI+10% serum at 37°C for 4 hr) of the parental and *efg1*ΔΔ strains derived from SC5314 and the indicated clinical isolates. Bars indicate mean of 4-5 fields from replicate experiments with error bars indicating standard deviation. B. Representative images of in vitro filamentation for the parental and *efg1*ΔΔ derivative of P57055. C. Comparison of the in vivo filamentation (24 hr post-infection) of the parental and *efg1*ΔΔ strains derived from SC5314 and the indicated clinical isolates. Bars indicate mean of 4-5 fields from replicate experiments with error bars indicating standard deviation. D. Representative images of in vivo filamentation for the parental and *efg1*ΔΔ derivative of P57055.

The TF Brg1 also plays an important role in the regulation of filamentation through a feedback loop with Nrg1, a repressor of filamentation (19, 20). Huang et al. found that deletion of *BRG1* reduced filamentation in all isolates in vitro (5) and we observed similar results in vitro (Fig. 6A/B). In vivo, *brg1*ΔΔ mutants were uniformly deficient in filamentation by at least 5-fold relative to the parental strain (Fig. 6C/D). These observations indicate that the filamentation master regulator status of Efg1 and Brg1 TFs is retained during filamentation in vivo.

**Figure 6.**
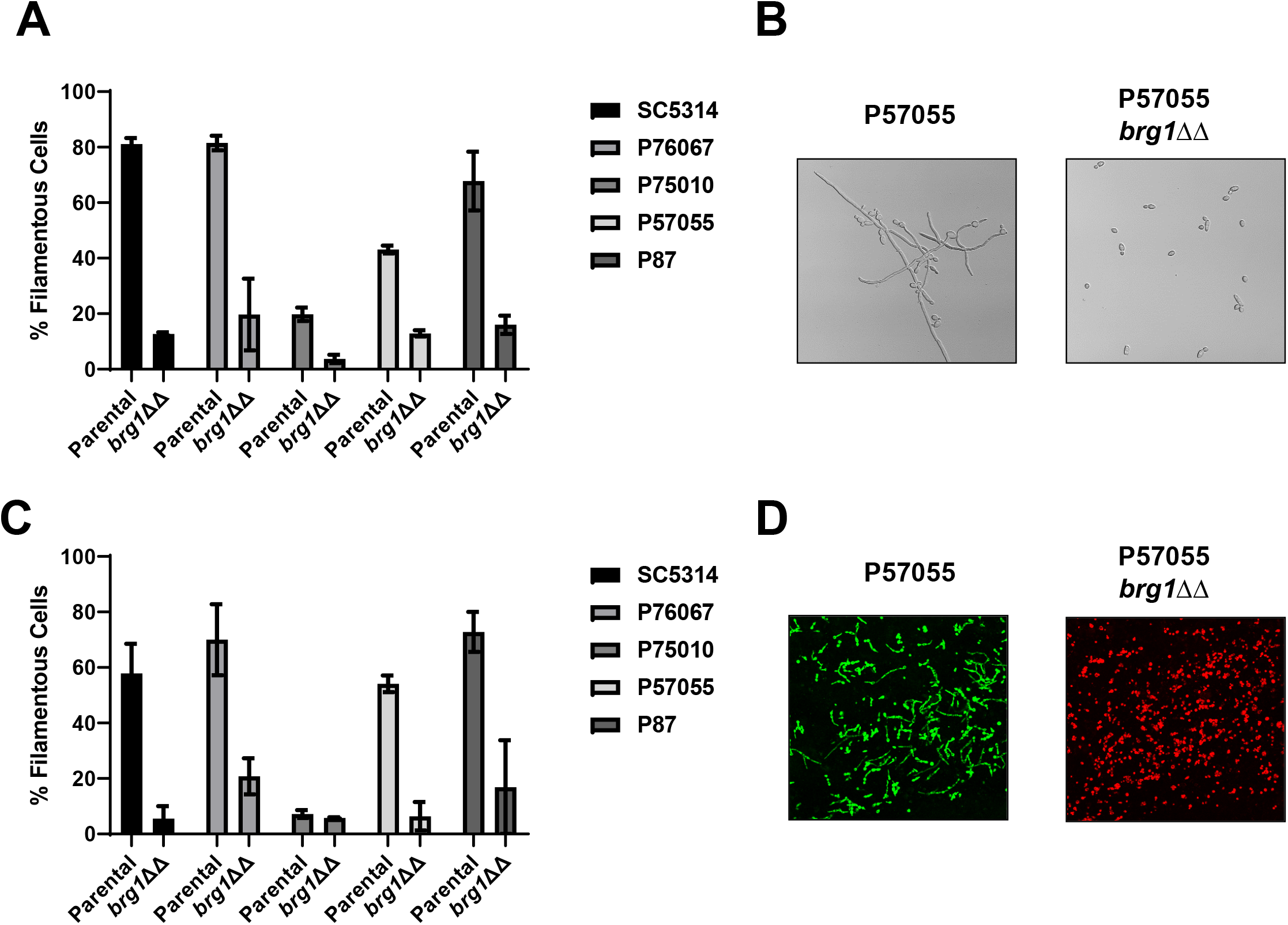
Brg1 is required for in vitro and in vivo filamentation across multiple *C. albicans* isolates. A. Comparison of the in vitro filamentation (RPMI+10% serum at 37°C for 4 hr) of the parental and *brg1*ΔΔ strains derived from SC5314 and the indicated clinical isolates. Bars indicate mean of 4-5 fields from replicate experiments with error bars indicating standard deviation. B. Representative images of in vitro filamentation for the parental and *brg1*ΔΔ derivative of P57055. C. Comparison of the in vivo filamentation (24 hr post-infection) of the parental and *brg1*ΔΔ strains derived from SC5314 and the indicated clinical isolates. Bars indicate mean of 4-5 fields from replicate experiments with error bars indicating standard deviation. D. Representative images of in vivo filamentation for the parental and *brg1*ΔΔ derivative of P57055.

### Ume6 plays a modest role during in vivo filamentation

In vitro, Ume6 is a well-characterized transcriptional regulator of filamentation whose expression has been shown to be necessary and sufficient to drive this process (21). Consistent with this role, deletion of *UME6* reduces filamentation 3-4-fold under in vitro conditions for all five strains (Fig. 7A/B). Under in vivo conditions, however, this level of reduction in filamentation was only seen in the *ume6*ΔΔ strain derived from P75010, the poorest filamenting strain (Fig. 7C/D). Deletion of *UME6* reduces filamentation by less than 1.5-fold for SC5314, P76067, and P87 while having a 2-fold effect on P57055; there was essentially no difference between the filamentation of P87 and its *ume6*ΔΔ mutant. Thus, most strains formed significant numbers of filaments in vivo in the absence of *UME6*. These observations indicate that the in vivo stimuli that lead to filamentation in vivo must trigger this process in a manner that largely bypasses the function of Ume6. Since Ume6 is required for filamentation under a variety of in vitro conditions (22), our data strongly support the hypothesis that the transcriptional networks for *C. albicans* vary with the specific environmental context.

**Figure 7.**
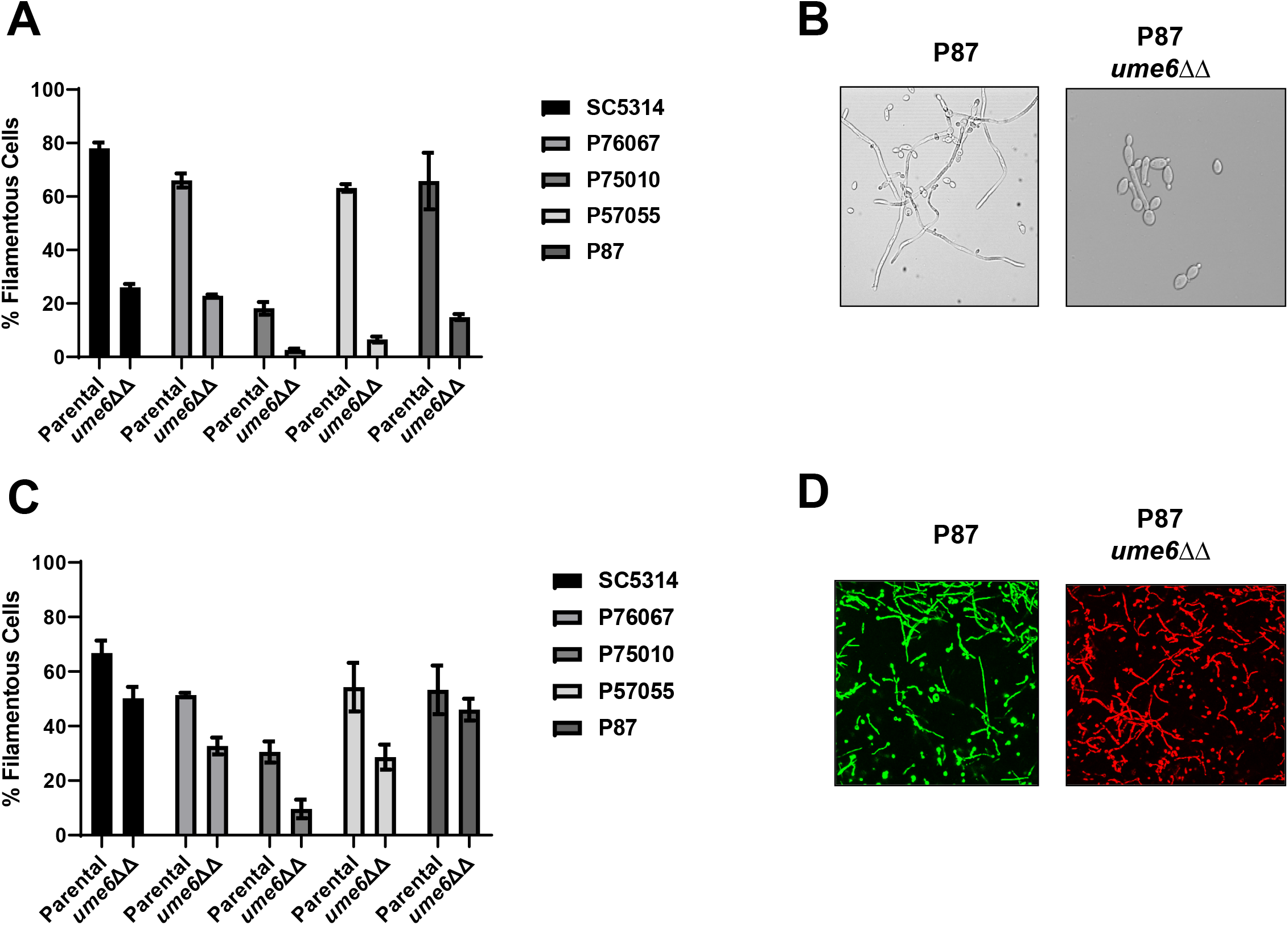
Ume6 has a more profound effect on filamentation in vitro than in vivo. A. Comparison of the in vitro filamentation (RPMI+10% serum at 37°C for 4 hr) of the parental and *ume6*ΔΔ strains derived from SC5314 and the indicated clinical isolates. Bars indicate mean of 4-5 fields from replicate experiments with error bars indicating standard deviation. B. Representative images of in vitro filamentation for the parental and *ume6*ΔΔ derivative of P87. C. Comparison of the in vivo filamentation (24 hr post-infection) of the parental and *ume6*ΔΔ strains derived from SC5314 and the indicated clinical isolates. Bars indicate mean of 4-5 fields from replicate experiments with error bars indicating standard deviation. D. Representative images of in vivo filamentation for the parental and *ume6*ΔΔ derivative of P87.

### Bcr1 is dispensable for filamentation in vivo

Bcr1 is a critical regulator of gene expression during biofilm formation both in vitro and in vivo (23, 24). Bcr1 has not typically been associated with the regulation of in vitro filamentation (4, 7), although it has been reported to negatively regulate the filamentation of opaque cells in vitro (25). Huang et al., however, found that Bcr1 regulated in vitro filamentation in P57055 and P87 but not in SC5314 or P76067 (5); our in vitro data matched those findings (Fig 8A/B). For strains for which in vitro filamentation was dependent on *BCR1*, Huang et al. also found that expression of *BRG1* was dependent on *BCR1* (5). In vivo, however, the deletion of *BCR1* has a minimal effect on the filamentation of any of the strains with the mutant forming filaments at a rate within 15% of the parental strain. Thus, in vivo filamentation is not dependent on the Bcr1-Brg1 interaction even in strains for which this regulatory circuit is required for filamentation in vitro. These observations further support the hypothesis that distinct transcriptional circuits regulate in vitro and in vivo filamentation.

**Figure 8.**
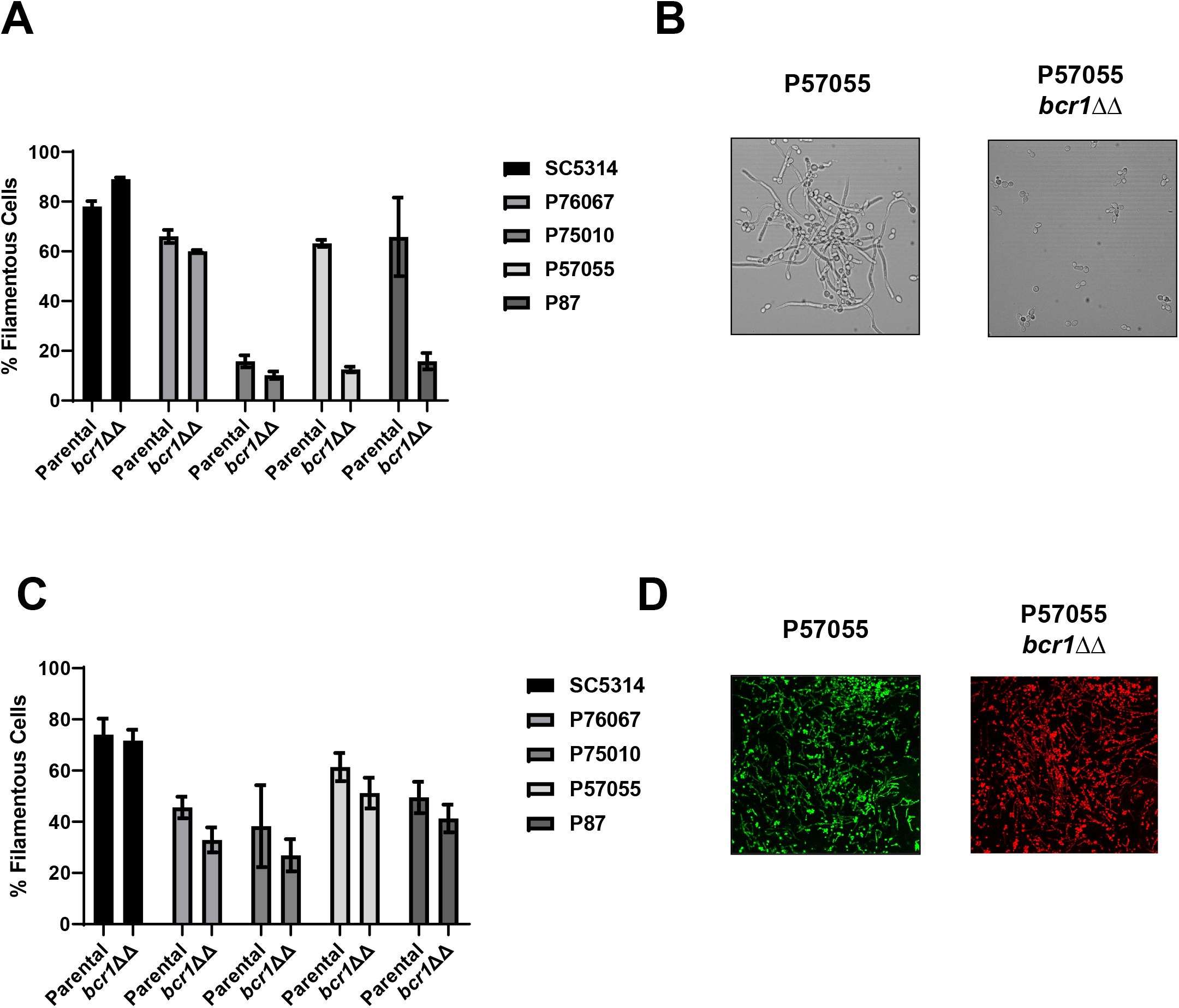
Bcr1 is dispensable for filamentation in vivo. A. Comparison of the in vitro filamentation (RPMI+10% serum at 37°C for 4 hr) of the parental and *bcr1*ΔΔ strains derived from SC5314 and the indicated clinical isolates. Bars indicate mean of 4-5 fields from replicate experiments with error bars indicating standard deviation. B. Representative images of in vitro filamentation for the parental and *bcr1*ΔΔ derivative of P57055. C. Comparison of the in vivo filamentation (24 hr postinfection) of the parental and *bcr1*ΔΔ strains derived from SC5314 and the indicated clinical isolates. Bars indicate mean of 4-5 fields from replicate experiments with error bars indicating standard deviation. D. Representative images of in vivo filamentation for the parental and *bcr1*ΔΔ derivative of P57055.

## Discussion

As the study of *C. albicans* pathogenesis matures, the application of new approaches to characterizing phenotypes and mechanisms during infection will provide more detailed models of the process required for infection and disease (9, 10). With this goal in mind, we have developed an intravital imaging strategy that allows the characterization of filamentation in an anatomic site that is relevant to the infection process. We have also taken advantage of the recent realization that the the study of clinical isolates with diverse ranges of phenotypes can provide important information not readily available by the study of domesticated laboratory reference strains (5, 13). These experiments have allowed us to make three major conclusions regarding the relationship between *C. albicans* filamentation in vitro and in vivo as discussed below.

Before we discuss these conclusions, it is important to consider how this model integrates with previous approaches to the assessment of *C. albicans* filamentation in vivo. For example, potential limitation of this approach is that the anatomic site of the infection may not be representative of other sites. However, our results for TFs such as *TEC1* and *UME6* correlate with previously reported kidney histology for these deletion strains (17, 22), suggesting there is significant overlap. However, it is important to note that *EFG1*, a canonical master regulator of filamentation, is not required for filamentation in the oral cavity of gnotobiotic pigs (26) and its deletion mutant is hyper-filamentous under in vitro embedded conditions (27). Furthermore, Witchley et al. have reported that *ume6*ΔΔ deletion mutants in the SN background filament similarly to WT cells in a model of commensal colonization of the GI tract (9). They also found that the *tec1*ΔΔ mutant was similar to wild type while *efg1*ΔΔ and *brg1*ΔΔ strains formed predominantly yeast. Our data are very similar to their findings indicating that Efg1 and Brg1 are important regulators of filamentation in multiple niches and that Ume6 plays a modest role in these niches. Thus, the phenotypes that we observe for these well-studied filamentation-related TFs are reasonably concordant with other examples of in vivo assessments of filamentation. The method of Witchley et al. (9) is a commensal counterpart to our approach in that both provide quantitative data in real time or near real time. One interpretation of our results in context with these other reports is that it seems likely that the transcriptional regulation of *C. albicans* filamentation varies from one niche to another. As such, genes required for filamentation in kidney, the most studied target organ in mice, may be different from the oral cavity or the submucosal stroma or the mucosa of the GI tract.

The major findings from our study include, first, by characterizing the in vivo filamentation of a laboratory reference strain (SC5314) and four clinical isolates with distinct virulence and in vitro filamentation phenotypes, we demonstrate that the relative abilities of the strains to filament in vivo correlate to some extent with in vitro filamentation in RPMI supplemented with 10% serum. This is summarized in Fig. 9 where the we have plotted the percentage of filamentous cells in vitro and in vivo for each parental strain and TF mutant examined in our study. The correlation between in vitro and in vivo filamentation was moderate (R^2^ = 0.607) with a slope less than 1; however, this correlation seems to be mainly driven by the high and low filamenting strains. Many of the mutants and strains had significantly discordant filamentation ratios when comparing in vitro and in vivo experiments.

**Figure 9.**
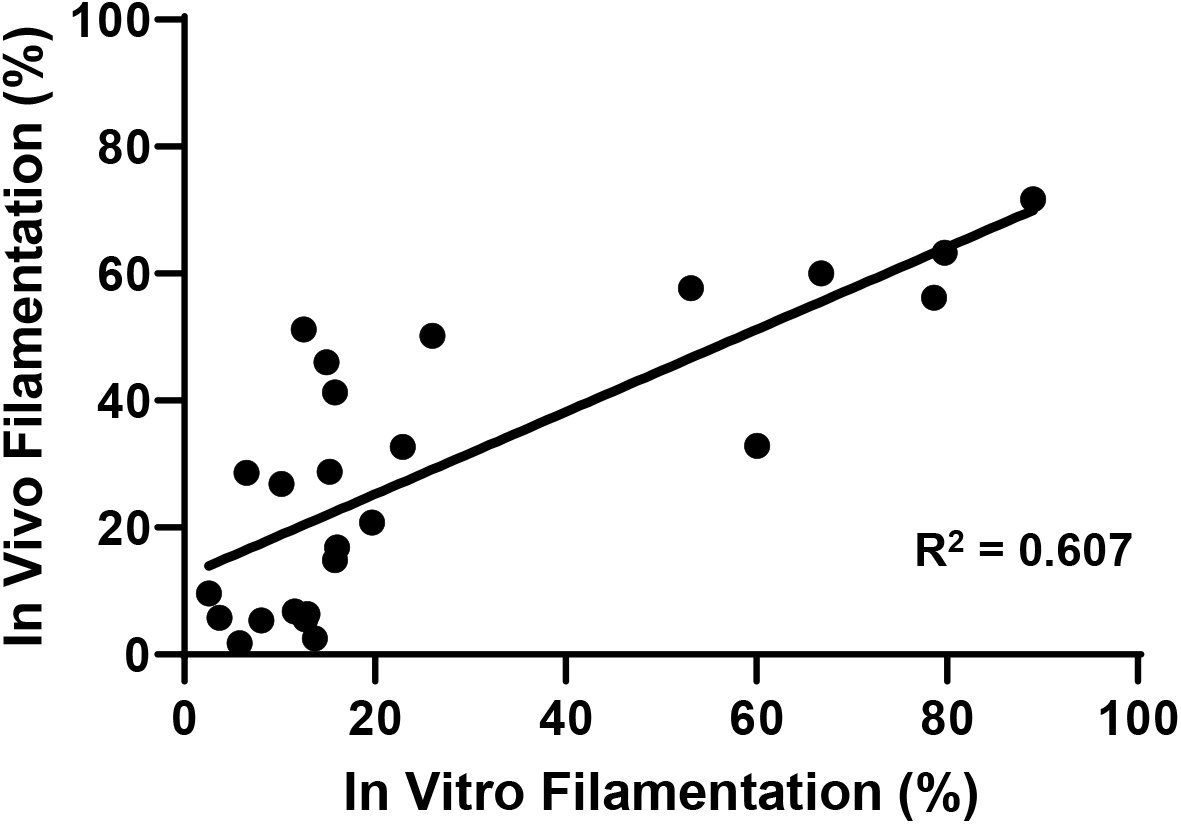
Summary comparison of in vitro and in vivo comparison of filamentation for each strain and mutant tested in this study. The mean % filamentation for each strain in vivo and in vitro is plotted. Linear regression was performed to assess the correlation between in vivo and in vitro filamentation for the set of strains. The R^2^ value is shown.

Although it is not possible to directly correlate the semi-quantitative plate-based, colony assays reported by Hirakawa et al. to our quantitative in vivo data (13), we found that one of the poorest filamenting strains (P57055) based on plate assays filamented quiet well in both RPMI+10% serum and in vivo. Not surprisingly, these results suggest that *C. albicans* is subject to signals that induce filamentation in vivo that are not replicated, either in type or extent, in vitro. Although not wholly unexpected based on recent studies demonstrating that *C. albicans* mutant strains vary in their in vitro filamentation phenotypes based on the specific induction conditions (6, 7), our experiments are the first to directly compare in vivo and in vitro filamentation phenotypes and make this observation. More strains will need to be examined to determine robustness of the general correlation between RPMI supplemented with serum and in vivo filamentation. It appears that *C. albicans* filamentation within colonies on agar plates is relatively distinct from that occurring within mammalian tissue (6).

We note that Tucey et al. recently reported filamentation data for the same set of strains during ex vivo infection of bone marrow-derived macrophages (28). Consistent with our in vivo and in vitro studies, P87 and SC5314 showed a high hyphal index. P75010 essentially formed no hyphae under these conditions while P76067 and P57055 also had low hyphal indices (28). The worst and best filamenting strains correlated well with our in vivo studies while the intermediate strains filamented much better in vivo than during macrophage infection. Tucey et al. found that strongly filamenting strains triggered NLRP3-mediated pyroptosis while those strains with lower hyphal indices did not (28). It would be interesting to be able to determine if these strains filament more robustly in macrophages within the host. Taken together, it appears that some strains of *C. albicans* form filaments robustly under most inducing condition while others require more specific conditions. Consequently, one should be cautious when making very general or absolute statements regarding the role of a strain or mutant in filamentation based observations made in vitro or ex vivo, particularly if only a few conditions are tested.

Our second major finding was that we found that the least virulent strain, P87, was able to filament strongly in vivo and in vitro. We also observed that the variation in in vivo filamentation amongst the five strains we studied was much less than the variation in virulence (14). Hirakawa et al. were unable to correlate in vitro filamentation with the virulence of these strain (13) and our data extend that lack of correlation to in vivo filamentation as well. In vitro, this lack of correlation was driven by the fact that seemingly non-filamentous strains such as P75010 are nonetheless virulent to a considerable degree (14). Our data suggest that this lack of correlation may be due instead to to the fact that strains with very different virulence phenotypes all form a significant number of filaments in vivo (at least 25-30%). Previously, Noble et al. reported that in vitro filamentation and the ability to establish infection were not well correlated in large-scale pooled infectivity screens (29). As such, our data provide additional support for the notion factors beyond filamentation are likely to contribute to the ability of *C. albicans* cause disease and that these factors vary in expression among clinical isolates.

Our third major finding is that, by studying a set of TF deletion mutants in the different *C. albicans* clinical strains, we have found that the function of specific TFs and TF circuits vary between in vivo and in vitro filamentation. Although Efg1 and Brg1 appear to retain their key roles regulating filamentation in vitro and in vivo, our observations suggest that the role of Ume6 is relatively modest in vivo. Under in vitro conditions, our data were consistent with previous reports that *UME6* deletion mutants have significant filamentation defects (21, 22). In the strongly filamenting strains SC5314 and P87, *ume6*ΔΔ mutants, the number and general quality of the filaments appeared to be quite similar to the parental strains. Indeed, the extent of filamentation observed for *ume6*ΔΔ mutants of SC5314 and P87 is very similar to histological sections of mouse kidney infected with a strain in which expression of *UME6* was transcriptionally repressed (22). The correlation of the *ume6*ΔΔ phenotypes with kidney histology further validates the ear infection model as representative of in vivo *C. albicans* filamentation. Overexpression of *UME6* increases filamentation and increases virulence and is clearly important for in vitro filamentation. However, it appears in vivo signals that stimulate filamentation do so in a manner that is largely Ume6-independent. We also found that the Bcr1-Brg1 circuit which is critical for in vitro filamentation and biofilm formation in some clinical strains (5) was not operative in vivo. These observations provide strong evidence that, although *C. albicans* filamentation is central part of its pathobiology, the TFs and transcriptional networks that regulate filamentation vary with the specific environmental cues that even critical regulators of this process in vitro can be bypassed in vivo.

Taken together, our data provide strong support for the notion that there are diverse regulatory mechanisms behind the complex phenotype of *C. albicans* filamentation and that these mechanisms vary with the specific niche or environmental context.

## Materials and Methods

### Strains, cultivation conditions, and media

The *C. albicans* clinical isolate strains and their respective mutants as well as the SN background-derived TF deletion mutants have been described previously (5, 7). All *Candida albicans* strains were pre-cultured over-night in yeast peptone dextrose (YPD) medium at 30°C. Standard recipes were used to prepare media (30). RPMI medium was purchased, supplemented with bovine serum (10% v/v).

### Strain construction

Fluorescently labelled strains were generated by using pENO1-NEON-NAT1 and pENO1-iRFP-NAT1 plasmid (8, 31). All transcription factor mutants were tagged with iRFP and their respective parent strains were tagged with green fluorescent protein (GFP). Briefly, the plasmids were digested with NotI enzyme for 2 hr at 37°C and subsequently, the linearized plasmid was further inserted into the *ENO1* locus (8). The *C. albicans* transformation was performed using the standard lithium acetate transformation method (32) and the transformants were selected using nourseothricin resistance marker (200 μg /ml, NAT; Werner Bioagents, Jena, Germany).

### Preparation and inoculation of mice with *C. albicans*

The mutant and their respective parent strains were grown overnight in YPD at 30°C. Harvested cells were washed thrice with sterile phosphate buffer saline (PBS) and counted with a hemocytometer. A 1:1 mixture of GFP-tagged reference strain and iRFP-tagged mutant strain was mixed to get a final count of 1× 10^8^ CFU/ml in PBS. A 5-6 weeks old, female DBA2/N mice (Envigo) used in these experiments were maintained on chlorophyll-free chow to minimize endogenous fluorescence. Prior to injections, the mice were anesthetized with isoflurane using SomnoSuite low flow anaesthesia machine (Kent Scientific) and the hair on the ears was removed by chemical depilation. 1 × 10^6^ CFU/ml (10 μl) of *C. albicans* cells containing equal volume of reference and mutant strains (1:1) were injected into the dorsal ear dermis of anesthetized mice with 29G1/2 needle. A characteristic papule was observed at the site of injection, indicating a successful intradermal injection.

### Confocal fluorescence microscopy

At 24 h post-injection, mice were anesthetized using isoflurane. Further mice were placed on the stage in the supine posture permitting ventral side of the ear facing downward for the imaging (10). Confocal images were carried out with a multiphoton laser scanning microscope (SP8; Leica Microsystem). The EGFP and iRFP was excited at 488nm and 635nm, respectively and emission was detected using a 505 to 525 nm and 655 to 755nm bandpass filter, respectively. The minimum of a 30 Z-stacks with an interslice interval between 0.57 μm was acquired with a 25X water immersive objective lens. The collected images were further max stacked using ImageJ software and used for analysis.

### Scoring criteria

We observed both pseudohyphal and hyphal cells under *in vivo* condition and for simplicity we termed them “non-yeast cells” whereas round and/ budded cells were termed as “yeast cells”. Yeast and non-yeast cells were quantified manually by following hyphae through each Z stacks (*n* => 100 cells). Statistical significance was determined by the unpaired Student’s t test.

### *In vitro* hyphal induction

C. albicans strains were incubated overnight in YPD at 30°C, harvested, and diluted into RPMI + 10% serum at 1:50 ratio and incubated at 37° C for 4 hours. Cells were collected and examined by light microscopy directly.

## Acknowledgements

The authors thank Rob Wheeler (Maine) for providing plasmids. We also thank Robb Cramer (Dartmouth) and Scott Filler (UCLA) for helpful comments on early versions of this manuscript. This work was supported by NIH grant 1R01AI33409 (DJK) and 5R01AI146103 (APM); the funders had no role in study design, data collection and interpretation, or the decision to submit the work for publication.

